# Corporal Punishment in Schools in the Jaffna District of Sri Lanka – A students’ view

**DOI:** 10.1101/2020.12.15.422836

**Authors:** M G Sathiadas, Annieston Antonyraja, Arunath Viswalingam, Shubanki Arunthavavinajagamoorthy

## Abstract

**Purpose:** Corporal-punishment has been prevailing in Sri- Lanka despite the strict law against it. Objectives of this study was to find the prevalence of corporal punishment in schools and the knowledge, perceptions and actions in the children.

**Methods:** A community based cross sectional descriptive study done among school students from Jaffna district using a pretested questionnaire containing 40 questions. Multistage stratified proportionate cluster sampling was used to recruit the students. Scoring systems were used to determine the knowledge and perception of Corporal-punishment. Data was analyzed using SPSS version 20.

**Results:** A total of 1130 students were recruited with mean age of 17.58±0.5y. A total of 687(60.7%) said they received some form of corporal punishment at least once during the school term. Physical punishment (64.5%) was common followed by psychological (27.1%). Teachers (54.8%) were the main group involved in corporal-punishment. Mean score for the knowledge was 11.11 and a good knowledge score was seen in 37.3% of the students. Participants with good knowledge favored law against it (p value<0.05), take legal action (P value<0.05) and supported complete prohibition of it (P value<0.05). Nearly 2% (n=22) had good perception and 40.8% (n=461) had a more positive perception. Majority (86.3%) preferred an alternative way of discipline and felt it was detrimental to the future (89.5%).

**Conclusion:** Corporal-punishment was experienced by majority of the students in school and the students were aware that it was illegal and those with good knowledge were willing to take legal action.

## Introduction

Corporal punishment is a method to discipline a child where an adult deliberately inflicts pain upon a child in response to a child’s unacceptable behaviour and instil discipline. The long-term aim is to prevent the recurrence of the bad behaviour and be consistent with the adult’s expectations [1].

Corporal punishment (CP) in the name of “discipline” continues in homes, schools, childcare institutions and juvenile detention centres [2]. Legal protection is given to children to protect them from corporal punishment and to ensure human rights. Almost 95% of the world’s total child population live in countries which lack a legal system to protect children and 29.3% live in South Asia. Nearly 55.7% of children live in countries that have no law to protect children from corporal punishment and 50.0% of them live in South Asia [3].

Both children and adults believe that CP is an effective way of discipline [4]. Many studies have shown the opposite by showing the detrimental effects of CP [5]. Some of them are poor academic performance, low level of class participation, avoiding school or dropping out for fear of getting beaten, declining self-worth or self-esteem and fear of teachers and school [6,7]. Prospective and meta-analysis have concluded that spanking fails to change the child’s behaviour but can cause longterm damage [8]. Physically abused children have an increased chance of developing depression, become more aggressive, behave antisocially and develop anxiety [9]. A study done in Sri Lanka in 2008 also showed that parental corporal punishment is associated with psychological harm in children [10].

Physical punishment used by an authoritative figure teaches violence as a method of correction and this can hinder parent-child relationship and other relationships [11, 12]. CP is partly blamed as an aetiology for criminal violence in youth hence it must be addressed as a preventive step [13].

The United Nations declared universal prohibition of corporal punishment in 2006. [14]. Global Initiative to end corporal punishment has been helping to mobilise and support nations to change their laws and social attitudes [15]. Sri Lanka expressed its commitment to prohibit corporal punishment at home and schools in July 2006, which was reiterated in 2017. Even though the instruction was adopted by a government circular, it is been confirmed through the enactment of legislation [16, 17].

Reporting and maintaining data on corporal punishment is vital to initiate preventive strategies. CP is likely to be under-reported as the victims are afraid of repercussions [7]. Unconfirmed reports say that most of the schools still practice CP and conventionally believe that mild CP is acceptable and necessary to discipline children [18]. There are few studies done regarding the prevalence of CP but very few regarding the knowledge and perception among the school children. Knowledge regarding CP is vital to understand the importance and the long term detrimental effect [19].

School children and teachers should be aware of CP and should take measure to prevent it. We conducted this study among school children to see the knowledge, perception, and actions the students would take towards CP. The objective of this study was to identify the prevalence of CP, factors that influence CP, their knowledge, its long-term effect on their wellbeing and the perception towards CP.

## Materials and Methods

### Setting

A community based cross sectional descriptive study was done among 16-19y studying in both private and government schools in the Jaffna district, Sri Lanka. The study period was July to August 2018. The sample size was calculated using the expected proportion of children experiencing corporal punishment being 50% with 95% confidence interval. To cluster design effect (DE) was considered as 1.8; and a non-repose rate of 5%. [20].

### Sampling

Multistage stratified proportionate cluster sampling was done, and ten schools located in the district of Jaffna were selected using a recently updated list of secondary schools in the district of Jaffna with a probability proportionate to the size of student population. Within each selected school, five classes were selected as the clusters to cover the different streams. A classroom comprising 20-30 students was considered as a cluster. The entire class that was considered as a cluster was sampled.

### Study tool

An anonymous pre-tested and standardised self-administered questionnaire was used. A field test was conducted with 10 experts in the field of child abuse to measure the content validity. The questionnaire was modified as per the expert suggestions, and the modified version was used as the study tool.

To assess the knowledge, questions regarding types of corporal punishments, the reason for the punishment and the legal implications were checked. A set of 17-items which had correct and incorrect responses were used, and each correct response was awarded +1. A score ≥ 13 was considered as good knowledge.

The perception towards CP was checked with another 22 items which recorded the response on the likert scale and the responses were recorded as “strongly agree” or “agree”, or “somewhat agree” or “disagree”, or “strongly disagree”. Depending on whether it was a proper attitude or not, scores from 1 to 5 were allotted. Reverse scoring was awarded for the negative responses. A score of ‘0’ was given for “don’t know”/ “can’t say”. A score of >85 was the considered as an overall good perception, 60-85 considered a more positive perception and <60 considered as needing a change in the perception.

The questionnaire also checked the preferred method of discipline and the outcome of CP in those who received it.

### Data analysis

Data was analysed by using Statistical Package for Social Sciences (SPSS) version 22.0. Responses were expressed as percentage and Chi-square test for significance of difference among proportions was calculated. Analysis of variance for significance of difference among means was calculated and Cronbach alpha was used to assess the reliability of the scores. The data was described using frequencies and percentages. One-way ANOVA was used to compare the means of the scores for perception and knowledge. P value of <0.05 was considered as statistically significant.

Ethical approval was obtained from the Ethics Review Committee of the University of Jaffna (*J/ERC/16/72/NDR/0143*) and permission was sought from the Provincial Educational authorities.

## Results

A total population of 1130 was recruited to the study. Majority were boys (53.5%) and mean age was 17.58±0.5y. (Table 1) Nearly 60.7% (687) had experienced at least one episode of punishment either at home or school during a school term. Physical type was experienced by 64.5% (n=443) followed by psychological seen in 27.1% (n=186) and both forms in 8.4% (n=58). Table 2 describes the number of incidences, the reason for CP and type of CP the child received. There was no significant relationship between age, sex, stream of study, family type, number of siblings and socio-economic status to the incidences of CP (p value>0.05). The students who received CP reported that the teachers (86.9%) were the main group inflicting CP followed by parents (74%), elder siblings (28%) and other elders in the community (10.3%).

**Table 1:** Socio demography of the study population.

| Item | Sub category | Number (%) |
| --- | --- | --- |
| **Mean age** | Girls (n=525) | 17.6±0.5yrs |
|  | Boys (n=605) | 17.5±0.5yrs |
| **Stream of study** | Maths | 225 (19.9) |
|  | Science | 294 (26.0) |
|  | Arts | 316 (27.9) |
|  | Commerce | 213 (18.8) |
|  | Information technology | 82 (7.3) |
| **Religion** | Hindus | 870 (76.9) |
|  | Christians | 248 (21.9) |
|  | Muslims | 12 (1.0) |
| **Socio-economic status** | <LKR 15,000 | 321 (28.4) |
|  | LKR 15,000-30,000 | 585 (51.8) |
|  | >LKR 30,000 | 214 (18.9) |
|  | Do not Know | 10 (0.8) |
| **Number of children in Family** | Only child | 198 (17.5) |
|  | 2-3 children | 586 (51.9) |
|  | >3 children | 346 (30.6) |
| **Family Type** | Nuclear | 815 (72.1) |
|  | Extended | 233 (20.6) |
|  | Single parent | 82 (7.3) |

**Table 2:** Corporal punishment experienced by the students.

| Item | Subcategory | Number (%) |
| --- | --- | --- |
| Type of Punishment@ (n=687) | Physical | 443 (64.5) |
|  | Psychological | 186 (27.1) |
|  | Mixed | 58 (8.4) |
|  | None | 443 (39.2) |
| Frequency of experiencing | Daily | 79 (7.0) |
|  | 1-3 per week | 348 (30.8) |
|  | 1-3 per month | 268 (23.7) |
|  | Once per term | 409 (36.1) |
| Place of experiencing the CP | School | 389 (56.6) |
|  | Home | 68 (9.9) |
|  | Community | 41 (6.0) |
|  | School and home | 189 (27.5) |
| Reason for the CP (Incidences) | Academic related<sup>a</sup> | 348 (50.6) |
|  | Discipline related<sup>b</sup> | 386 (56.2) |
|  | Behaviour related<sup>c</sup> | 184 (26.8) |
|  | Reason not clear | 67 (9.8) |

<sup>a</sup>Academic related: making mistakes, not reading/writing, not doing home work, low marks
<sup>b</sup>Discipline related: coming late, absent, fighting with others, damage to property, being noisy,
<sup>c</sup>Behaviour related: stealing, not getting permission to go out, telling lies,

The common form of physical punishment was beating with a stick or hand experienced by 46.6% (n=527). The common form of psychological CP was being called names (20.8%) followed by shouted and yelled (19.8%). (Table 3)

**Table 3:** Prevalence of strategies of punishment.

| **Type of physical punishment** | **Number of incidences (%)** |
| --- | --- |
| Ear squeezing | 339 (30.0) |
| Slapping on the cheeks | 367 (32.5) |
| Asked to kneel-down | 340 (30.0) |
| Sit ups | 244 (21.6) |
| Beating with a stick/cane/hand to the lower limbs | 527 (46.6) |
| Pinching | 235 (20.8) |
| Stand on the table | 178 (15.8) |
| Standing on one leg for a long time | 94 (8.3) |
| Hands-above-the head | 163 (14.4) |
| Stand outside the class/Stand for long time | 457 (40.4) |
| **Type of psychological punishment** | **Number (%)** |
| Shouted, yelled, and using language not acceptable | 224 (19.8) |
| Threatened to chase or spank | 145 (12.8) |
| Called out names like lazy, crazy, dumb, donkey, pig etc | 235 (20.8) |
| Compared with other students | 189 (16.7) |
| Listing out the mistakes in front of class | 175 (15.5) |

Some forms of punishments that the students experienced were perceived as favourable to them such as getting extra homework in 58% (n= 655), after school class in 38% (n=429), cleaning the school and grounds experienced by 23% (n=260), removal of privileges in 29% (n=328) and when the teacher/parent explained and gave advice in 47% (n=531)

Nearly 7.3% (n=82) 0f the children received medical treatment following an injury due to CP. Of the students who received CP, 60.4% (n=415) said it did not change or affect them, 18.6% (n=128) said it changed their behaviour and 9.0% (n=62) said it affected them psychologically as they felt sad and rejected.

The mean score for the knowledge component was 11.2±3.2. There was a significant difference in the mean score for knowledge between boys and girls. 85% of the students knew the child’s right charter and nearly all the children knew the emergency contact number to dial if a harm occurred to them. Table 4 describes the components of knowledge.

**Table 4:** Knowledge and perception regarding corporal punishment.

| Knowledge regarding corporal punishment |  |  |  |
| --- | --- | --- | --- |
|  | Girls (n=525) | Boys (n=605) | Statistical analysis |
| Mean score of knowledge | 11.43±3.1 | 10.9±3.5 | One way ANOVA<br>F(1,1130)=7.47<br>P=0.006 |
| Good Knowledge (scored >13) | 210 (40.0%) | 212 (35%) | Pearson’s X<sup>2</sup>(1,1130)=<br>2.95 P=0.08 |
| Knowledge factors* |  |  |  |
| Aware of the United nations charter | 446 (84.9%) | 514 (84.9%) | Pearson’s X<sup>2</sup>(4,5650)=<br>246.5 P<0.0001 |
| Know the emergency number to call | 504 (96%) | 558 (92.2%) |  |
| CP is Illegal | 397 (75.6%) | 457 (75.5%) |  |
| CP affects the future | 398 (75.8) | 415 (68.6%) |  |
| CP is a punishable offence | 402 (76.6%) | 426 (70.4%) |  |
| Perception regarding corporal punishment |  |  |  |
|  | Girls (n=525) | Boys (n=605) | Statistical analysis |
| Mean score of attitudes 54.9±14.2 | 54.9±13.4 | 54.9±15.5 | One-way ANOVA<br>(F1,1130) =0.01<br>P=0.9 |
| Classification of perception score |  |  |  |
| Overall good feeling (>85) | 10 (1.9%) | 12 (2.0%) | X<sup>2</sup>(2,1130) =0.01<br>P=0.9 |
| More towards a positive perception (60-85) | 214 (40.7%) | 247 (40.8%) |  |
| Change in perception is needed (<60) | 301 (57.3%) | 346 (57.1%) |  |

\*Cronbach alpha was 0.75

The perception of CP indicated a mean population score of 54.9±14.2. The scores were not statistically significant between the girls and boys. Nearly 2% (n=22) had good perception, 40.8% (n=461) had a more positive perception and 57.3% (n=647) needed a perceptual change towards CP. (Table 4)

A total of 38.2% (n=432) of the participants agree while 27.1% (n=306) disagree that CP promotes future perpetrators to commit child abuse. Nearly 70.4% (n=796) of the participants disagreed with the statement “CP is psychologically beneficial” and 45.9% (n=519) agreed that corporal punishment is associated with an increase in delinquent and antisocial behaviours in childhood. It was noted that students marked either strongly agree or agree in 60.7% (n=686) to “CP disgraces you”, 57.9%(n=654) to “CP causes too much pain and has a detrimental effect”, 45.6% (n=515) to “increases the antisocial and illegal behaviour”, 56.7% (n=641) to aggressive behaviour and 38.3% (n=50) to increases conflicts in the family and leads to gender based violence. The mean scores of each component tested for the study population is indicated in table 5.

**Table 5:**
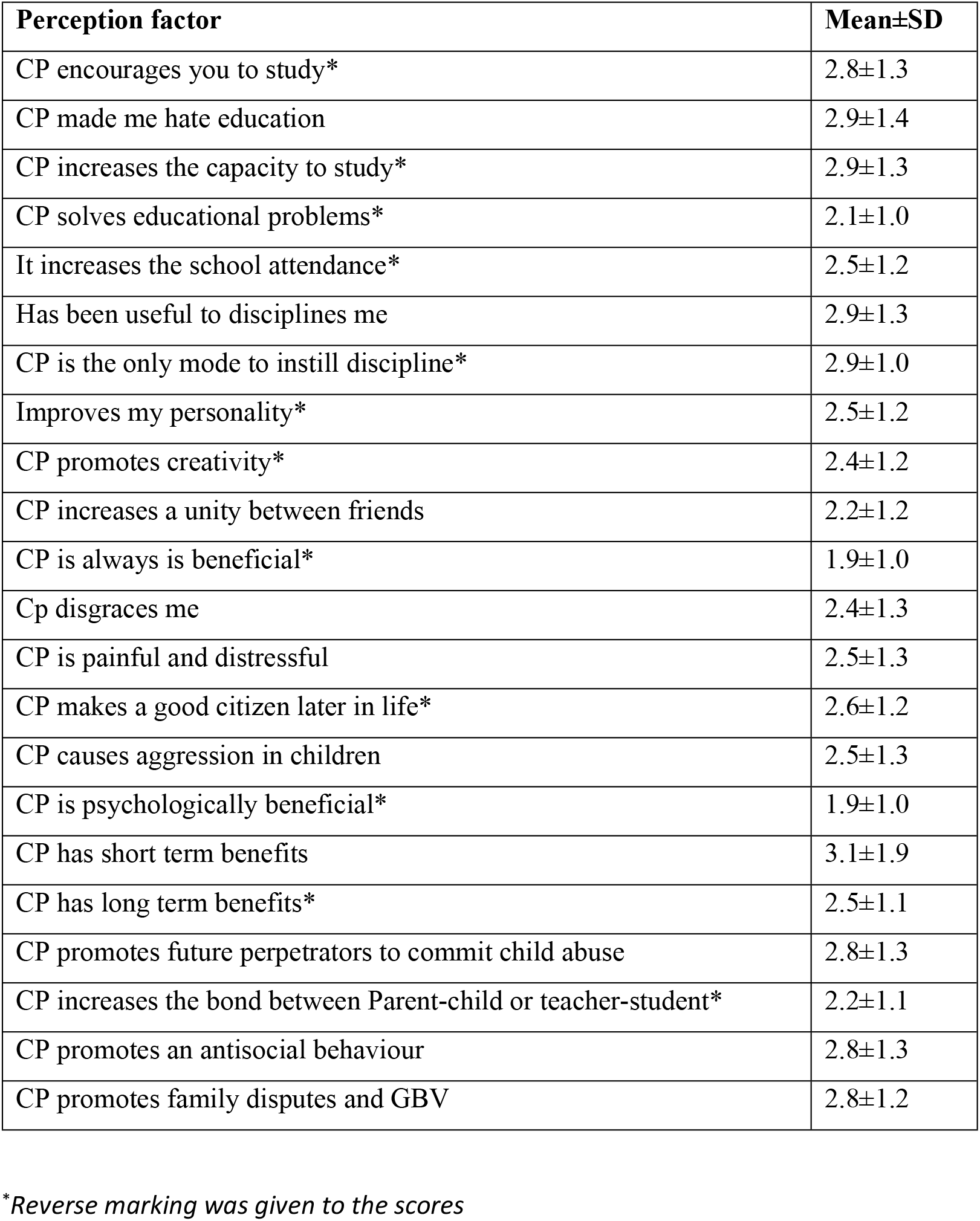
Mean Scores of perception of corporal punishment.

Majority (86.3%) preferred an alternative way of discipline and felt corporal punishment had a detrimental effect on their future development (89.5%). A total of 31.8% (n=359) said it was a method used to discipline students and accepted it with reluctance. Nearly 79.6% (n=900) answered yes to “will you take legal actions against CP” and 50.5% (n=571) supported complete prohibition of corporal punishment.

Participants with good knowledge and positive perception favor legal action, support complete prohibition of CP and felt it had a detrimental effect on their future. (Table 6)

**Table 6:**
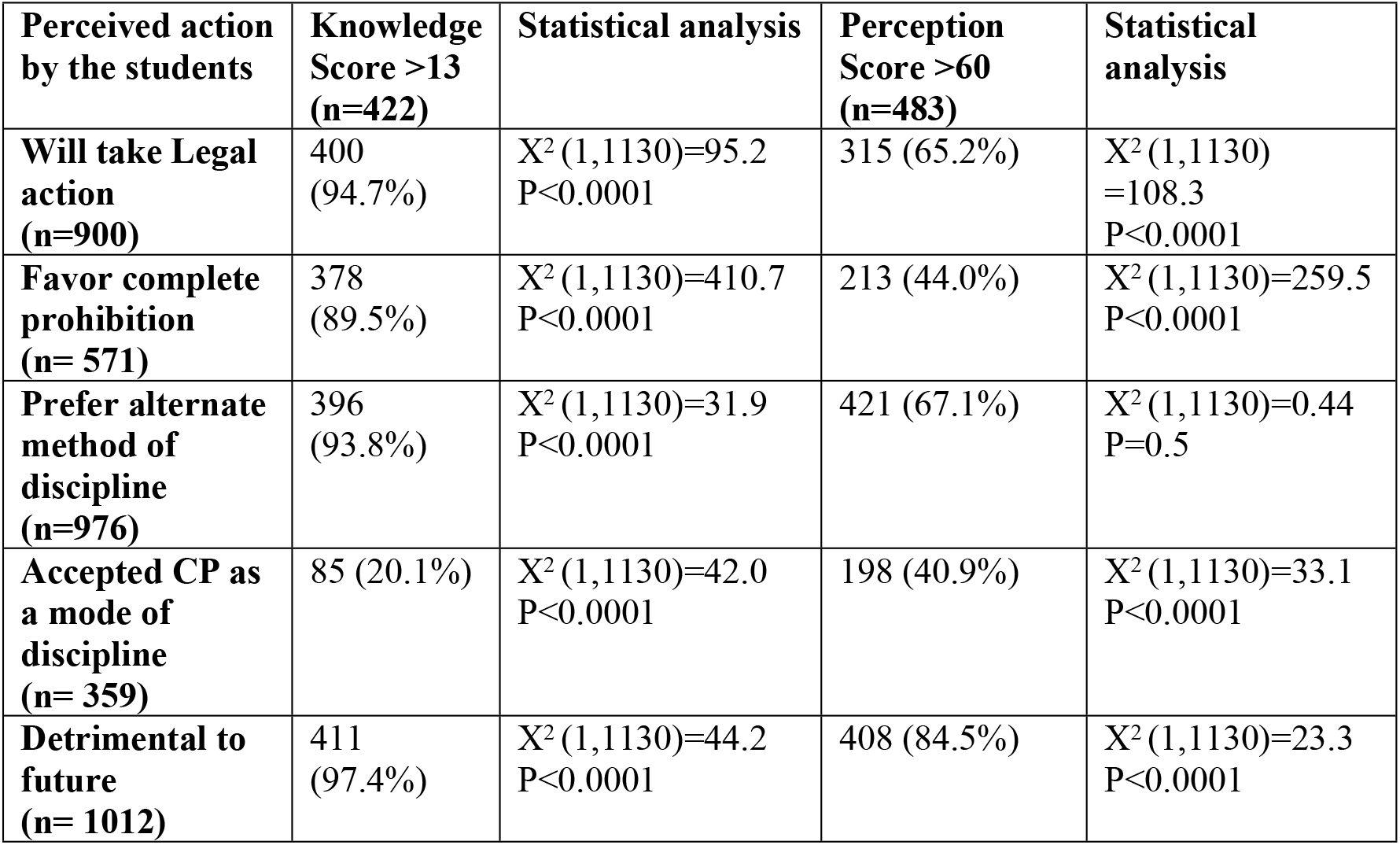
Comparative analysis of perceived action to knowledge and perception scores.

| **Perceived action by the students** | **Knowledge Score >13 (n=422)** | **Statistical analysis** | **Perception Score >60 (n=483)** | **Statistical analysis** |
| --- | --- | --- | --- | --- |
| **Will take Legal action (n=900)** | 400 (94.7%) | $X^2(1,1130)=95.2$<br>$P<0.0001$ | 315 (65.2%) | $X^2(1,1130)=108.3$<br>$P<0.0001$ |
| **Favor complete prohibition (n= 571)** | 378 (89.5%) | $X^2(1,1130)=410.7$<br>$P<0.0001$ | 213 (44.0%) | $X^2(1,1130)=259.5$<br>$P<0.0001$ |
| **Prefer alternate method of discipline (n=976)** | 396 (93.8%) | $X^2(1,1130)=31.9$<br>$P<0.0001$ | 421 (67.1%) | $X^2(1,1130)=0.44$<br>$P=0.5$ |
| **Accepted CP as a mode of discipline (n= 359)** | 85 (20.1%) | $X^2(1,1130)=42.0$<br>$P<0.0001$ | 198 (40.9%) | $X^2(1,1130)=33.1$<br>$P<0.0001$ |
| **Detrimental to future (n= 1012)** | 411 (97.4%) | $X^2(1,1130)=44.2$<br>$P<0.0001$ | 408 (84.5%) | $X^2(1,1130)=23.3$<br>$P<0.0001$ |

## Discussion

Teachers and parents use various tactics to discipline and correct behaviour in children. Despite having legal implications against corporal punishment, schools in the northern part still use it to discipline children. This hinders the long-term developments and is known to cause psychological problems and may lead to criminal behaviour in adult life [9]. Sri Lankan schools thrive to show off their educational and extra-curricular performances. This leads to immense stress in the teachers and parents and thereby it is transmitted to the children. This stress and anxious situation make the adults resort to physically and psychologically harmful methods to make the children perform [21]. In this study also physical and psychological punishments have been predominant when compared to positive methods of discipline. Nearly 65% have reported having the experience of CP in our study which is also similar to a study from India which showed 65% and a study from Sri Lanka stated nearly 80% have experienced it [7, 20]. Studies done in India have demonstrated that CP commonly occurs in poor socio-economic background children, but our study did not demonstrate this [3, 7].

The reason for CP is mainly physical seen in 64.5% (n=443) of the population and when compared to a previous study done in 2017 the incidence was nearly 80%. The awareness created on litigation may have contributed to the reduction in the prevalence from 2017 to our study. The downward trend is promising as the Millennial developmental goal is to end violence against children by 2030 [14].

CP is known to cause negative effect like physical injuries, poor academic performances, school dropouts, low self-esteem and fear of school and teachers. In our study the short-term harm of physical injury was seen in 7.3%. Despite these effects CP is still advocated by the children and adults. Our study also showed that 31.8% (n=359) accepted CP despite the knowledge that it had a detrimental effect.

Nearly 36% of the students have experienced CP at least once during the school term and 7% have experienced it daily. Of the various modes most common one was beating with the stick or cane experienced by 46.6% of the students followed by standing outside the class for long time (40.4%), slapping the cheek (32.5%) and ear squeezing (30%). Similar types of injuries have been seen in other studies [7, 20]. Most of the children encountered the punishments in school but 27.5% experienced CP both at home and school. Similar results were seen in other studies [8, 20].

Psychological aggression towards the children was noted in 27.1% and calling by names was the commonest (20.8%). When compared to other studies this seems to be low and it may be that the students’ perception of psychological trauma may be different in different study populations and countries [22].

The knowledge regarding CP was good in 40% of the girls and 35% of the boys. The easy access to social media and web may have contributed to the good knowledge and girls generally being more disciplined were better aware of CP than the boys [23].

The perception showed mixed responses as the scoring system showed good and more positive feeling only in 42.7%. and nearly 57% needed a perceptual change. 608(53.8%) of the students strongly agreed or agreed to corporal punishment disgraces them. Almost equal percentage of participants said that CP made them to hate studies (47.3%) and CP encourages them to learn (45.8%). This may be supported by the cultural belief that the teacher or the parent does this for their betterment [24]. Violence in this culture is accepted and thereby CP becomes acceptable to many. CP leads to higher levels of societal violence, thereby reducing use of corporal punishment should lead to reductions in societal violence. This awareness may help to abolish CP in the future [24].

The action to overcome CP was promising as most (86.4%) preferred an alternative method of discipline despite 31.8% accepting CP as a mode of discipline. Nearly 89% felt it was detrimental to their future. Even though most indicated CP was detrimental and supported complete prohibition they accepted it as a mode to discipline. This may be a cultural impact where the children are taught from early years that the adults inflict harm to correct unacceptable behaviour and to discipline them. [24].

Students with good knowledge and a positive attitude regarding CP, had a clearer plan of action. This shows a promising outlook towards the future.

To shape desirable behaviour in students and correct misbehaviour disciplinary strategies rather than punishment strategies are required [25]. The power dynamics of adults and the cultural acceptability of CP must be abolished, and alternative approaches must be encouraged. Some of the alternative approaches are non-violent discipline, effective communication and conflict resolution [26]. When Positive discipline is inculcated it promotes appropriate behaviour. Teachers should be made aware of alternatives to corporal punishments in schools to stop the menace of punishing children and making the schools safe [27].

This study is limited to16-18-y-old adolescents and cannot be generalized as younger age groups were not considered. A similar study in teachers and parents will give a better idea regarding the perception which will abolish CP completely.

## Conclusion

Use of corporal punishment is widespread with physical harm being common. The knowledge regarding CP was satisfactory but a change in perception is needed among the school students attending secondary schools in the Northern Province.

## DECLARATION

### Ethics approval

Ethical approval was obtained from the Ethics Review Committee of the Faculty of Medicine, University of Jaffna (*J/ERC/16/72/NDR/0143*) and permission was sought from the Provincial Educational ministry and relevant zonal directors.

### Consent to participate

Eligible sample of participants were informed and written informed consent from the parent or guardian and accent from the students were obtained. Participants received instructions for opting out of the survey.

### Consent for Publication

Not applicable in this study

### Availability of data and materials

The data in this study is available from the corresponding author on request.

### Code availability

Not applicable to this study

### Author contribution

MGS - Designed and developed the protocol, monitored data collection, reviewed and revised the manuscript and approved the final document, AA - Data collection, analysis and approved the final manuscript, AV – Data Collection, manuscript preparation and approved the final manuscript, SA - Data collection, manuscript preparation and approved the final manuscript, All Authors read and approved the final manuscript prior to submission.

## List of Abbreviations

CP: Corporal Punishment
WHO: World Health Organisation
GCE A/L: General Certificate of Education Advanced Level
UNICEF: United Nations Childrens’ Fund

